# Spared cognitive and social function following perinatal ablation of ATRX despite transient microglia dysregulation

**DOI:** 10.1101/2025.08.06.668920

**Authors:** Kasha Y. Mansour, Sarfraz Shafiq, Nathalie G. Bérubé

## Abstract

Mutations in the chromatin-remodelling factor ATRX underlie syndromic ID and ASD, yet the contribution of microglial ATRX to early brain development remains unknown. We used a tamoxifen-inducible Cx3cr1-CreERT2 system to delete *Atrx* in microglia during the first postnatal week, achieving >90% recombination at one month of age. At both one and three months, ATRX-null microglia displayed a sustained increase in CD68-positive phagocytic foci and in Ki67+ proliferative microglia, but an unexpected ∼20% reduction in microglial density, despite only modest evidence of caspase-3-mediated apoptosis. A comprehensive behavioural battery conducted between three and six months revealed normal anxiety, locomotion, learning, memory, social interaction and sensory gating, coinciding with a decline in knockout efficiency to ∼45% in cortex and ∼75% in hippocampus, suggestive of microglial repopulation by ATRX-expressing cells. These findings demonstrate that perinatal loss of ATRX drives chronic microglial reactivity and turnover with no discernable effects on neurobehaviours, suggesting a developmental resilience of neural circuits to transient microglial dysregulation.

## Introduction

The X-linked *ATRX* gene encodes a SWI/SNF-family chromatin-remodelling ATPase that orchestrates transcriptional regulation and preserves genomic integrity through interactions with multiple protein partners [1]. Hypomorphic variants in *ATRX* cause ATR-X syndrome, a syndromic form of intellectual disability that can also present with autistic traits, α-thalassemia, microcephaly, seizures, and a spectrum of craniofacial, genital, and skeletal anomalies. Pathogenic *ATRX* mutations have also been reported in cases of ASD [2]. Because *ATRX* resides on the X chromosome, hemizygous males are predominantly affected, while most heterozygous females are protected by skewed X-chromosome inactivation that preferentially silences the mutant allele [3], [4].

Microglia are highly dynamic cells that are commonly referred to as the immune cells of the central nervous system. In their resting state, microglia act to maintain homeostasis of the brain by supporting the neuronal networks, as well as by surveying for changes in the microenvironment and attending to any damage [5]. In response to various pathologies, microglia can adopt different states characterized by transcriptomic and proteomic signatures [6] as well as undergo morphological changes and release various cytokines, depending on the stimulus encountered [7]. Importantly, microglia play key roles during brain development by shaping the neural circuitry. They can simultaneously release factors that promote neuronal survival and initiate programmed cell death, maintaining a delicate balance necessary to ensure the correct number of neurons [8], [9]. They are responsible for removing excess synapses that are generated during development to ensure proper connectivity [10]. Microglia also regulate myelination through phagocytosis of oligodendrocyte progenitor cells during development [10], [11]. Impaired synaptic pruning and abnormal myelination have been associated with neurodevelopmental disorders, including intellectual disability and autism spectrum disorder [11], [12].

In previous work, we conditionally deleted *Atrx* in microglia at postnatal day 45 and found that the loss of ATRX caused a reactive phenotype: microglia showed elevated CD68 foci and underwent a transient burst of proliferation. These changes coincided with altered morphology and electrophysiological properties of hippocampal CA1 pyramidal neurons and produced measurable deficits in long-term spatial memory [13].

Given that pathogenic *ATRX* variants in humans cause ID and ASD, and that microglia exert their strongest influence on circuit formation during early development, we now ask whether removing ATRX from microglia in the perinatal period elicits equivalent or more severe deficits. We deleted ATRX selectively in microglia through administration of tamoxifen to lactating dams at postnatal days P2-P4, triggering CreERT2-mediated recombination in their pups. One month later, immunofluorescence staining confirmed >90% ATRX knockout in microglia and revealed a reactive phenotype typified by increased CD68 staining, increased Ki67+ nuclei, and a 20% decrease in microglia density. Despite these cellular changes, male mice tested between 3-6 months of age were indistinguishable from controls in assays for anxiety, locomotion, learning, memory, social interaction, repetitive behaviour and sensorimotor gating. Follow-up experiments at three months revealed that a much lower proportion of microglia lacked ATRX, suggesting partial repopulation by wildtype cells.

## Results

### Compromised homeostasis of ATRX-null microglia

Perinatal microglia-specific *Atrx* deletion was achieved with the tamoxifen-inducible Cx3cr1-CreERT2 driver line. Female *Atrx* floxed mice were mated with Cx3cr1-CreERT2 males, and nursing dams received tamoxifen intraperitoneally (1 mg/day) on postnatal days 2–4. Pups ingested the drug via maternal milk, activating CreERT2 exclusively in microglia and excising the floxed *Atrx* allele, thereby generating ATRX microglial knockout (ATRX mi-KO) offspring (**Fig. 1a**). As ATR-X syndrome primarily affects males, only the male pups were used in our experiments. Immunofluorescence analysis of one-month-old brain sections confirmed efficient and cell-type-restricted deletion: 85–95% of Iba1⁺ microglia in the cortex and hippocampal subfield lacked ATRX immunoreactivity, while surrounding neurons and astrocytes retained ATRX expression (**Fig. 1b,c**).To test whether the loss of ATRX alters microglial homeostasis, we quantified phagocytic activity in coronal brain cryosections from one-month-old mice that were double-labelled for CD68, a lysosomal/phagocytic marker, and Iba1, a pan-microglial marker. Across all animals examined (n = 20 cells, N = 3 brains each genotype), the ATRX mi-KO group displayed a robust increase in the number of CD68-positive puncta per Iba1-positive cell in every region analyzed. One-way ANOVA revealed significant elevations in the cortex (F_2,6_ = 34.71, p = 0.0005), CA1 (F_2,6_ = 9.53, p = 0.0137), CA2 (F_2,6_ = 24.51, p = 0.0013), CA3 (F_2,6_ = 19.73, p = 0.0023) and dentate gyrus (F_2,6_ = 34.13, p = 0.0005) when compared with both Cre-negative and Cre-positive controls (**Fig. 2a,b**). Given that the Cx3cr1-CreERT2 driver alone was reported to trigger mild microglial activation [14], we directly compared the two control cohorts. Aside from a modest rise in CD68 burden within the dentate gyrus of Cre-positive brains (p = 0.0373), no significant differences were detected in the cortex (p = 0.3265), CA1 (p = 0.2962), CA2 (p = 0.1088) or CA3 (p = 0.7585) (**Fig. 2a,b**). These findings indicate that increased CD68 seen in ATRX-deficient microglia is attributable to the loss of ATRX itself rather than to Cre expression.

**Figure 1.**
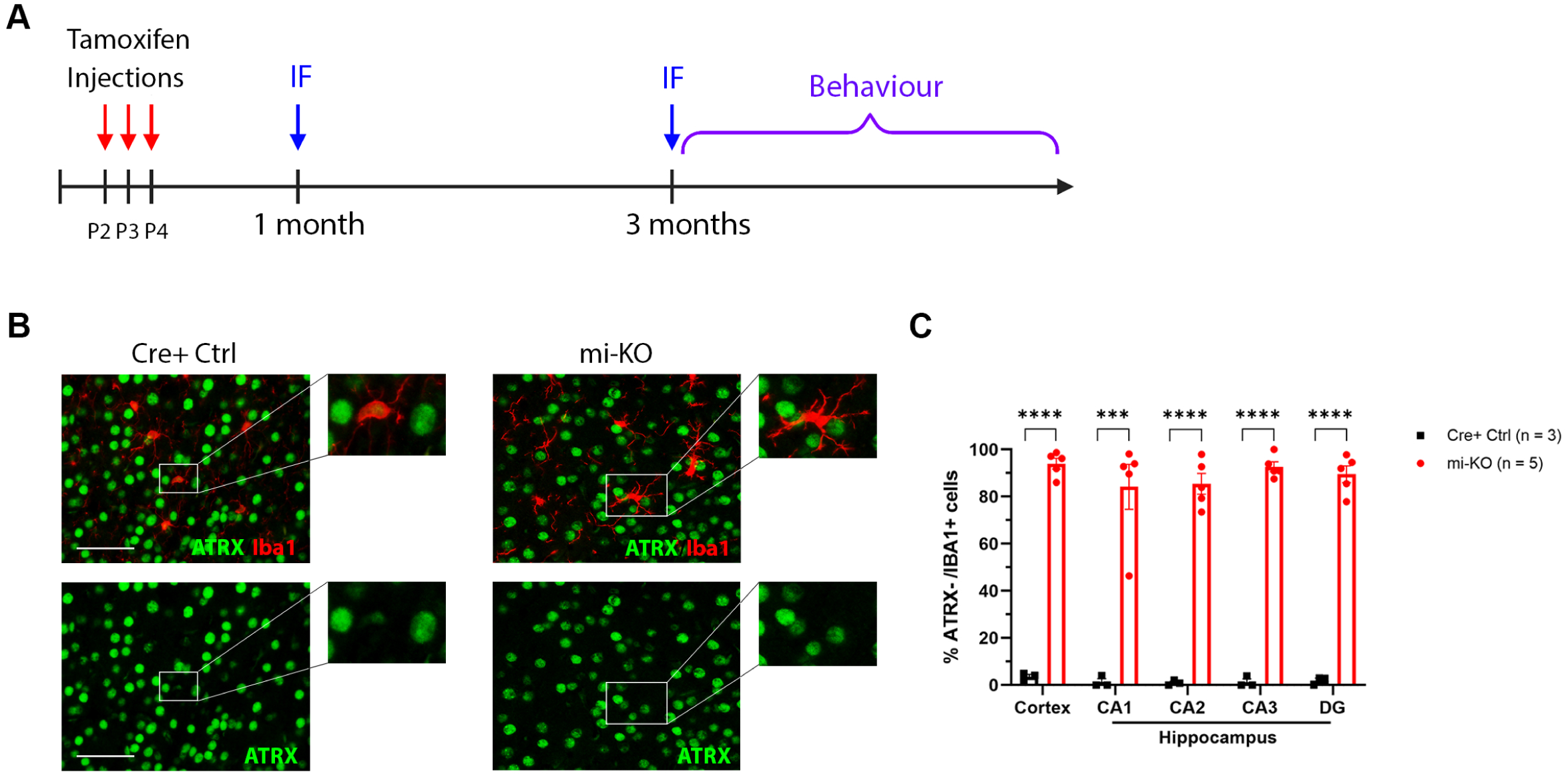
Conditional deletion of ATRX in microglia is >90% efficient one month after tamoxifen induction. **(a)** Experimental design. Lactating dams received intraperitoneal tamoxifen injections on post-natal days (P) 2–P4 (red arrows). Brains were collected for immunofluorescence at 1 and 3 months of age (blue arrow), and behavioural assays were performed between 3 and 6 months (purple bracket). **(b)** Representative confocal images of coronal brain cryosections stained for ATRX (green) and the microglial marker Iba1 (magenta); nuclei are counter-stained with DAPI (grey). Scale bar, 50 µm. **(c)** Quantification of microglial ATRX loss. Bars show the percentage of Iba1+ cells that are ATRX negative in Cre+ controls (Ctrl) and ATRX mi-KO; the number of mice (n) is indicated in parentheses. Data are mean ± SEM; unpaired two-tailed Student’s t-test. ***P < 0.001; ****P < 0.0001.

**Figure 2.**
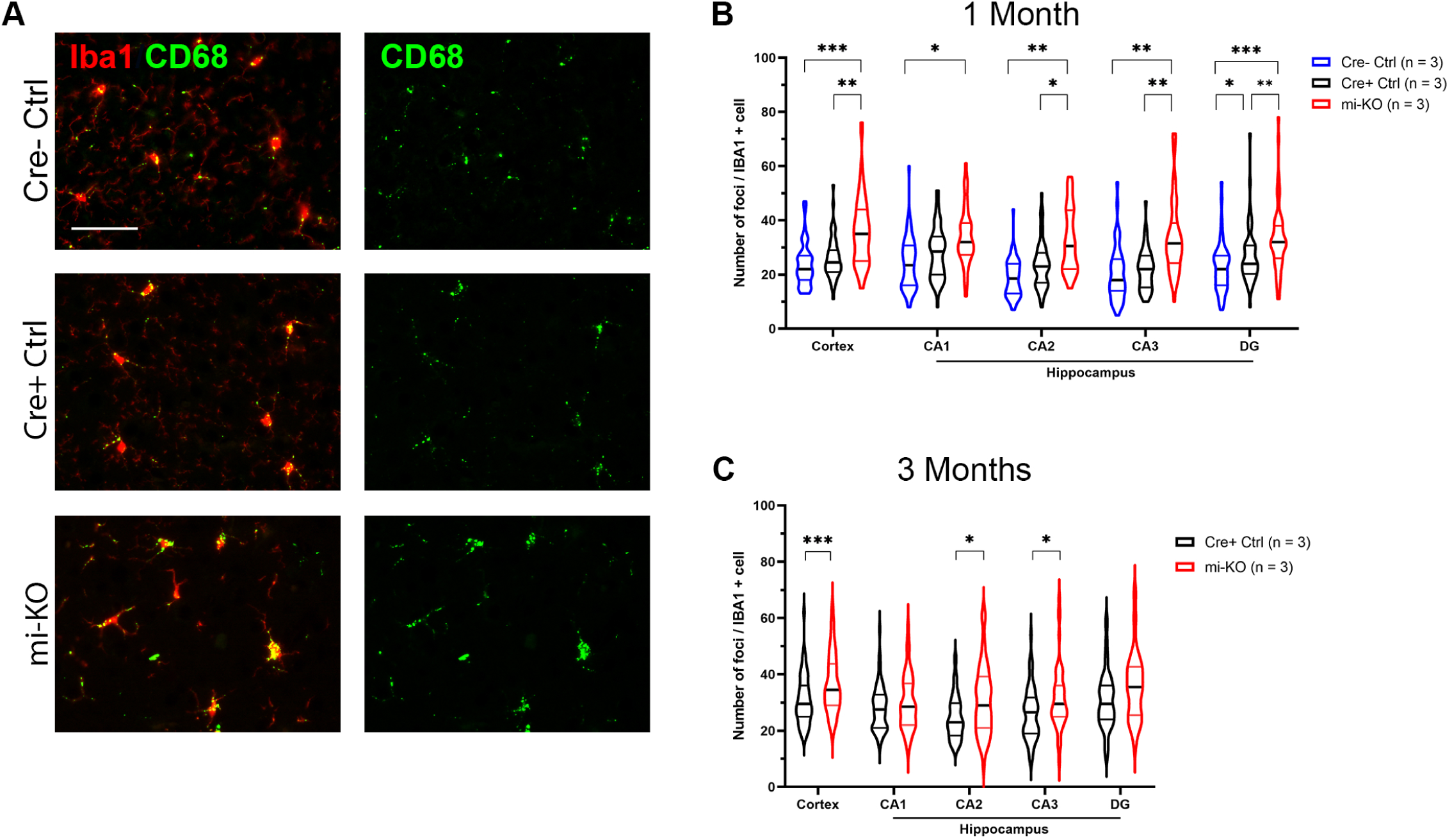
Ablation of ATRX in microglia leads to increased CD68 immunoreactivity. **(a)** Immunofluorescence staining of cortical cryosections of 1 month old brains with CD68 (a marker of phagocytic microglia) and the microglial marker IBA1. Scale bar represents 50 µm. Number of CD68+ foci per Iba1+ cell at **(b)** 1 month (one-way ANOVA) and **(c)** 3 months (t-test). Cre- Ctrl: Cre- control, Cre+ Ctrl: Cre+ control, mi-KO: ATRX microglia knockout. The number (n) of mice used for each genotype is indicated in parentheses. Data is presented as the median, and the 25th to the 75th percentile of values are indicated. * = P<0.05, ** = P<0.01, *** = P<0.001.

To assess the longevity of this phenotype, we repeated the analysis in three-month-old mice. Elevated CD68 staining per microglia persisted in the cortex (p = 0.0004) and in the CA2 (p = 0.0406) and CA3 (p = 0.0432) regions of the hippocampus, whereas CA1 (p = 0.3431) and dentate gyrus (p = 0.1414) values had returned to control levels (**Fig. 2c**). Together, these data demonstrate that microglia lacking ATRX adopt a chronic reactive state that is most pronounced in the cortex and selected hippocampal subfields.

### Increased proliferation yet decreased density of microglia in the ATRX miKO brains

To determine whether ATRX loss alters microglial cell-cycle dynamics, we analyzed coronal sections from one-month-old ATRXmi-KO;Ai14 reporter mice and their littermate controls. Sections were co-stained for Ki67, a canonical proliferation marker, and imaged in parallel with Ai14, which selectively labels Cre-recombined microglia. The proportion of Ki67⁺/Ai14⁺ cells was significantly higher in ATRX-deficient brains than in either Cre-positive or Cre-negative controls, with robust increases detected in the cortex (p = 0.0049) and in the CA1 (p = 0.0007), CA2/3 (p = 0.0013) and dentate gyrus (p = 0.0001) subfields of the hippocampus (N = 3; **Fig. 3a,b)**. These data indicate that ATRX-null microglia enter the cell cycle more frequently, a finding consistent with the heightened phagocytic activity documented earlier.

**Figure 3.**
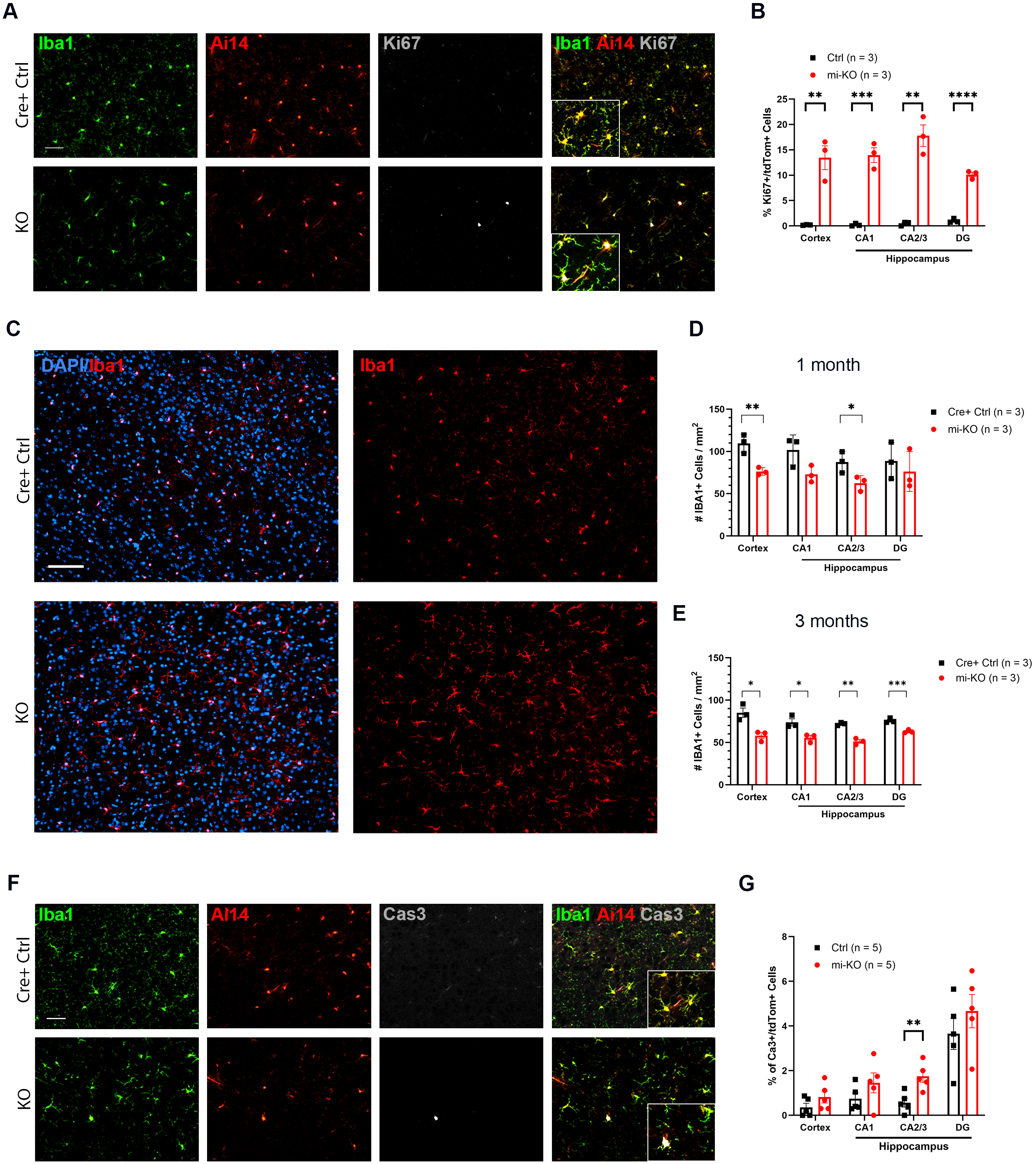
Reduced microglia density despite proliferative burst in ATRX miKO cortex and hippocampus. **(a)** Immunofluorescence staining of brain cryosections from 1 month old mice expressing the Ai14 Cre reporter with Iba1 and the proliferative marker Ki67. **(b)** Quantification of the percentage of tdTom+ that are Ki67+ in the cortex and hippocampal regions (t-test). **(c)** Immunofluorescence staining of brain cryosections from 1 month old mice with the microglial marker Iba1. Quantification of Iba1+ cell density at **(d)** 1 month and **(e)** 3 months (t-test). Ctrl: Cre+ control, mi-KO: ATRX microglia knockout. **(f)** Immunofluorescence staining of brain cryosections from 1 month old mice expressing the Ai14 Cre reporter with Iba1 and the apoptosis marker cleaved caspase 3 (Cas3). **(g)** Percentage tdTom+ that are Cas3+ in the cortex and hippocampal regions (t-test). The number (n) of mice used for each genotype is indicated in parentheses. Data is presented as mean ± SEM. * = P<0.05, ** = P<0.01, *** = P<0.001, **** = P<0.0001.

Surprisingly, this proliferative surge did not translate into a larger microglial population. Quantification of Iba1 immunoreactivity revealed a significant reduction in microglial density in the cortex and CA2/3 of ATRX miKO brains, with a similar downward trend in CA1 and the dentate gyrus (cortex p = 0.0084; CA2/3 p = 0.0481; CA1 p = 0.0701; DG p = 0.5363; N = 3; **Fig. 3c,d**). When the analysis was repeated in three-month-old animals, microglial density was diminished in all regions examined (cortex p = 0.0146; CA1 p = 0.0232; CA2/3 p = 0.0010; DG p = 0.0027; **Fig. 3e**), indicating that the discrepancy between proliferation and cell number widens with age.

A plausible explanation for the reduced cellularity is an increase in microglial death. We therefore stained adjacent sections for cleaved caspase-3. ATRX-deficient brains showed a modest elevation in Caspase-3⁺/Ai14⁺ cells that reached statistical significance in the CA2/3 region (p = 0.0072) but not in the cortex, CA1 or dentate gyrus (p = 0.1887, 0.2041 and 0.3522, respectively; N = 5; **Fig. 3f,g**). Although this partial rise in apoptosis may contribute to the loss of microglia, additional mechanisms are likely required to fully account for the marked reduction in cell density.

### Ablation of ATRX does not perturb oligodendrocyte-lineage homeostasis or motor performance

During early postnatal development microglia eliminate surplus oligodendrocyte progenitor cells (OPCs) [10], [11]. Given the hyper-phagocytic phenotype of ATRX-deficient microglia, we asked whether OPC abundance is altered in these mice. Coronal sections from one-month-old brains were immunolabelled for platelet-derived growth factor receptor-α (PDGFRα), a selective OPC marker. Quantitative analysis showed no difference in OPC density between ATRX miKO and Cre-positive control mice in the cortex, CA1, CA2/3, or dentate gyrus (N = 3 brains per genotype; cortex p = 0.386, CA1 p = 0.169, CA2/3 p = 0.554, DG p = 0.424; **Fig. 4a,b**). Since OPC differentiation and subsequent myelination influence locomotor coordination, we evaluated motor learning on the accelerating rotarod. Across ten six-minute acquisition trials, the latency to fall did not differ between ATRX miKO (n = 9) and control (n = 6) animals (F_1,13_ = 0.37, p = 0.55; **Fig. 4c**). When tested 24 h later for retention, both groups again performed equivalently over four trials (F_1,13_ = 0.49, p = 0.50; **Fig. 4d**). Together, these findings indicate that perinatal loss of ATRX in microglia neither disturbs OPC homeostasis nor impairs gross motor learning and memory.

**Figure 4.**
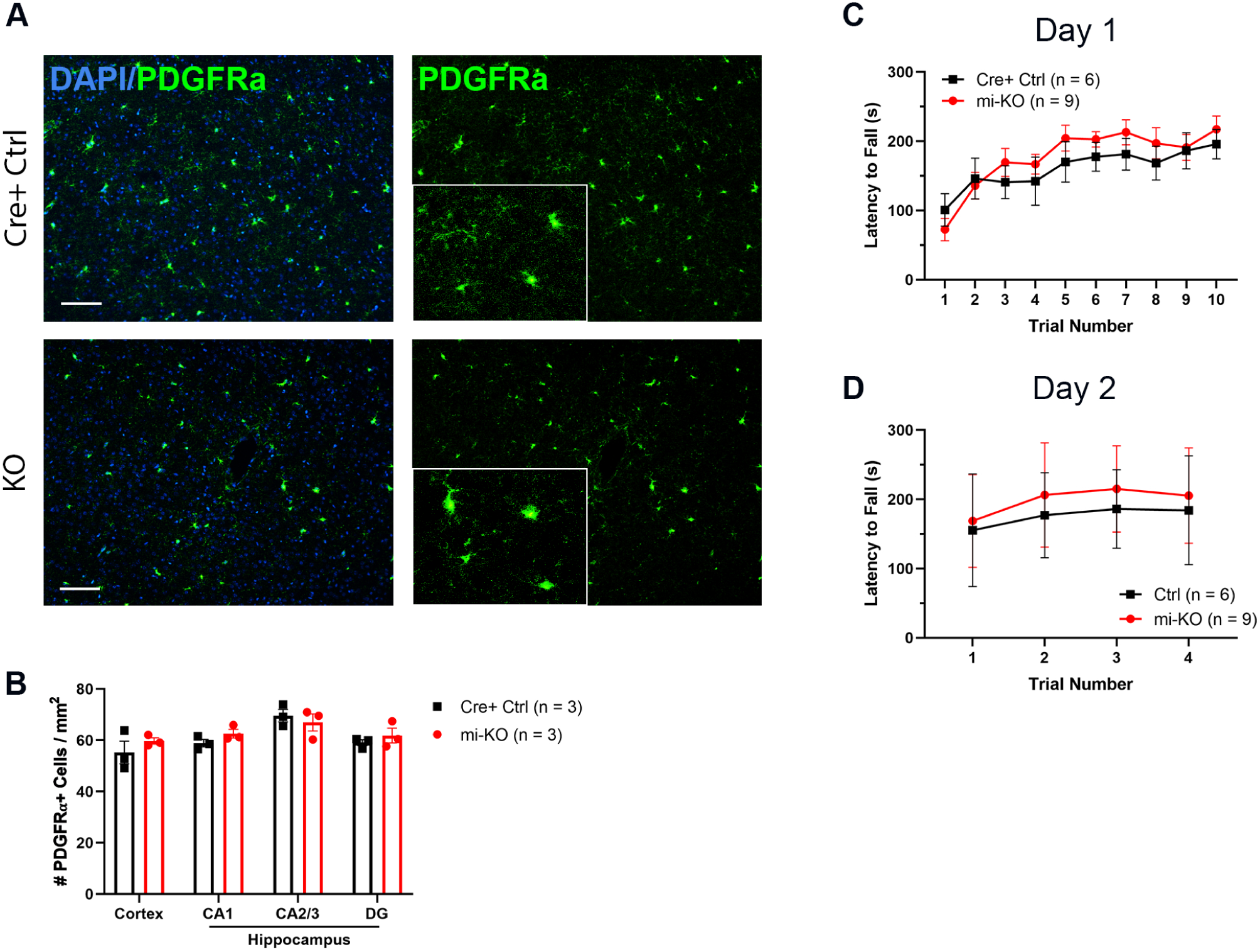
No changes in the density of oligodendrocyte progenitor cells, and no motor deficits caused by perinatal ATRX deletion in microglia. **(a)** Immunofluorescence staining of brain cryosections from 1 month old mice with PDGFRα, a marker of oligodendrocyte progenitor cells. Scale bar represents 50 µm. **(b)** The number of PDGFRα+ cells per square millimeter (t-test). **(c)** The latency to fall off the rotarod during 6-minute trials on day 1 and on **(d)** day 2 (two-way ANOVA). Ctrl: Cre+ control, mi-KO: ATRX microglia knockout. The number (n) of mice used for each genotype is indicated in parentheses. Data is presented as the mean ± the SEM.

### Microglial ATRX deletion does not alter anxiety-like behaviour or cognitive performance

To determine whether perinatal loss of ATRX in microglia influences emotional reactivity, we first evaluated anxiety-like behaviour in adult males. In the light–dark box, ATRX mi-KO mice spent the same proportion of time in the illuminated compartment as Cre-positive controls (Ctrl, n = 21; mi-KO, n = 28; two-tailed t-test, p = 0.78). No group differences emerged in the elevated- plus maze, where the percentage of time spent in the open arms was comparable between genotypes (Ctrl, n = 20; mi-KO, n = 28; p = 0.29). Similar results were obtained in the open-field test, in which the time spent in the central area did not differ significantly (Ctrl, n = 21; mi-KO, n = 27; p = 0.08; Fig. 5**a–c**). Collectively, these assays indicate that microglial ATRX ablation does not modify baseline anxiety levels.

**Figure 5.**
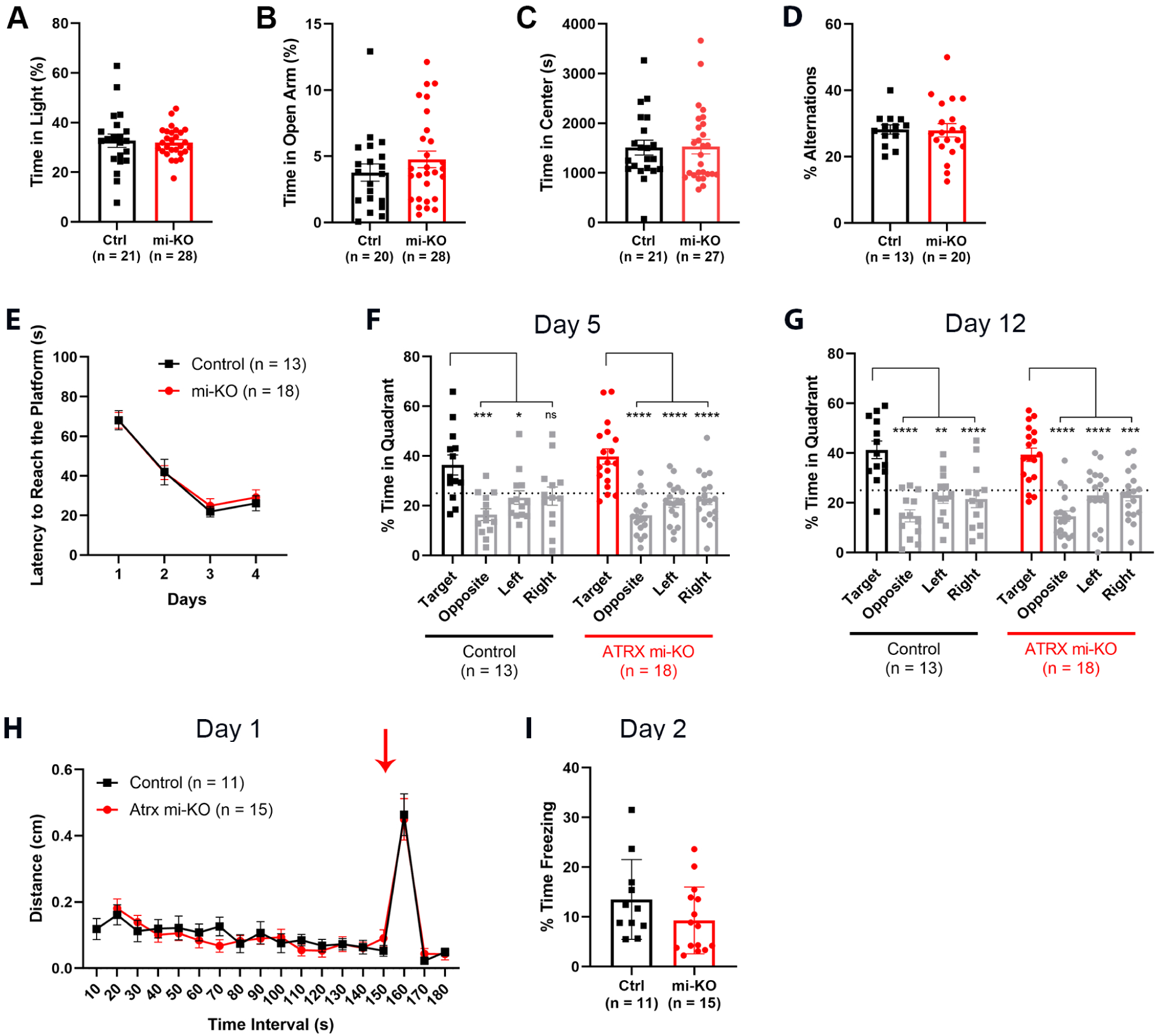
No changes in anxiety or memory deficits observed upon perinatal deletion of ATRX in microglia. **(a)** The amount of time spent in the light area during the light/dark box test (t-test), **(b)** the amount of time spent in the open arm of the elevated plus maze (t-test), and **(c)** the amount of time spent in the centre during the 2-hour open field test (t-test). **(d)** Percent alternations in the Y-maze. **(e)** Latency to reach the platform in the Morris water maze over 4 days of training (4 trials/day) (two-way ANOVA). Time spent in each quadrant on **(f)** day 5 and **(g)** day 12, the dotted line indicates 25% (two-way ANOVA). **(h)** Distance travelled over 10 second intervals on day 1 of the contextual fear conditioning task (two-way ANOVA), the arrow indicates the time at which the shock was delivered. **(i)** Total time spent freezing on day 2 (t-test). Ctrl: Cre+ control, mi-KO: ATRX microglia knockout. The number (n) of mice used for each genotype is indicated in parentheses. Data is presented as mean ± SEM. * = P<0.05, ** = P<0.01, *** = P<0.001, **** = P<0.0001.

Since *ATRX* mutations cause intellectual disability in humans, we next assessed several domains of learning and memory. Working memory, measured by spontaneous alternation in the Y-maze, was intact: both groups alternated at equivalent rates (Ctrl, n = 13; mi-KO, n = 20; p = 0.94; **Fig. 5d**). Spatial learning in the Morris water maze proceeded normally. During the four-day acquisition phase the latency to locate the hidden platform declined similarly in controls and ATRX mi-KO mice (Ctrl, n = 13; mi-KO, n = 18; repeated-measures ANOVA, genotype effect F₁,₂₉ = 0.15, p = 0.70; **Fig. 5e**). Probe trials on days 5 and 12 confirmed intact spatial memory: each genotype spent significantly more time in the target quadrant than in non-target quadrants (day 5, ATRX mi-KO F₃,₆₈ = 18.98, p < 0.0001; controls F₃,₄₈ = 6.50, p = 0.0009; day 12, ATRX mi-KO F₃,₆₈ = 17.63, p < 0.0001; controls F₃,₄₈ = 13.79, p < 0.0001; **Fig. 5f,g**). Associative memory was also unaffected: 24 h after foot-shock training, contextual fear conditioning elicited comparable freezing in ATRX mi-KO and control mice (Ctrl, n = 11; mi-KO, n = 16; p = 0.47; **Fig. 5h,i**).

Overall, despite noticeable cellular and phagocytic phenotypes, perinatal deletion of ATRX in microglia leaves adult anxiety-like behaviour, working memory, spatial learning, and associative memory intact.

### Microglial ATRX loss fails to elicit autism-relevant behavioural phenotypes

*ATRX* mutations are occasionally linked to autism spectrum disorder. We therefore asked whether perinatal deletion of ATRX in microglia reproduces core ASD-like traits in mice, namely perseverative behaviour, hyperactivity, altered sociability, and impaired sensory gating [2].

Repetitive behaviour was assessed in three complementary paradigms. During the 2-h open-field session the number of vertical rearing episodes was identical in ATRX mi-KO and Cre+ control animals (Ctrl, n = 21; mi-KO, n = 27; p = 0.93; **Fig. 6a**). Likewise, the percentage of marbles buried in a 30-min marble-burying task and the cumulative time spent grooming in an induced self-grooming assay were indistinguishable between genotypes (marble burying, p = 0.80; grooming, p = 0.89; **Fig. 6b,c**). These data indicate that ATRX-deficient microglia do not promote stereotypy. Total distance travelled in the same open-field arena, an index of locomotor drive, was comparable across the 2-h session (p = 0.19; **Fig. 6d**). Breaking the trace into 5-min bins uncovered no genotype effect (F₁,₄₆ = 1.74, p = 0.19; **Fig. 6e**), ruling out hyperactivity.

**Figure 6.**
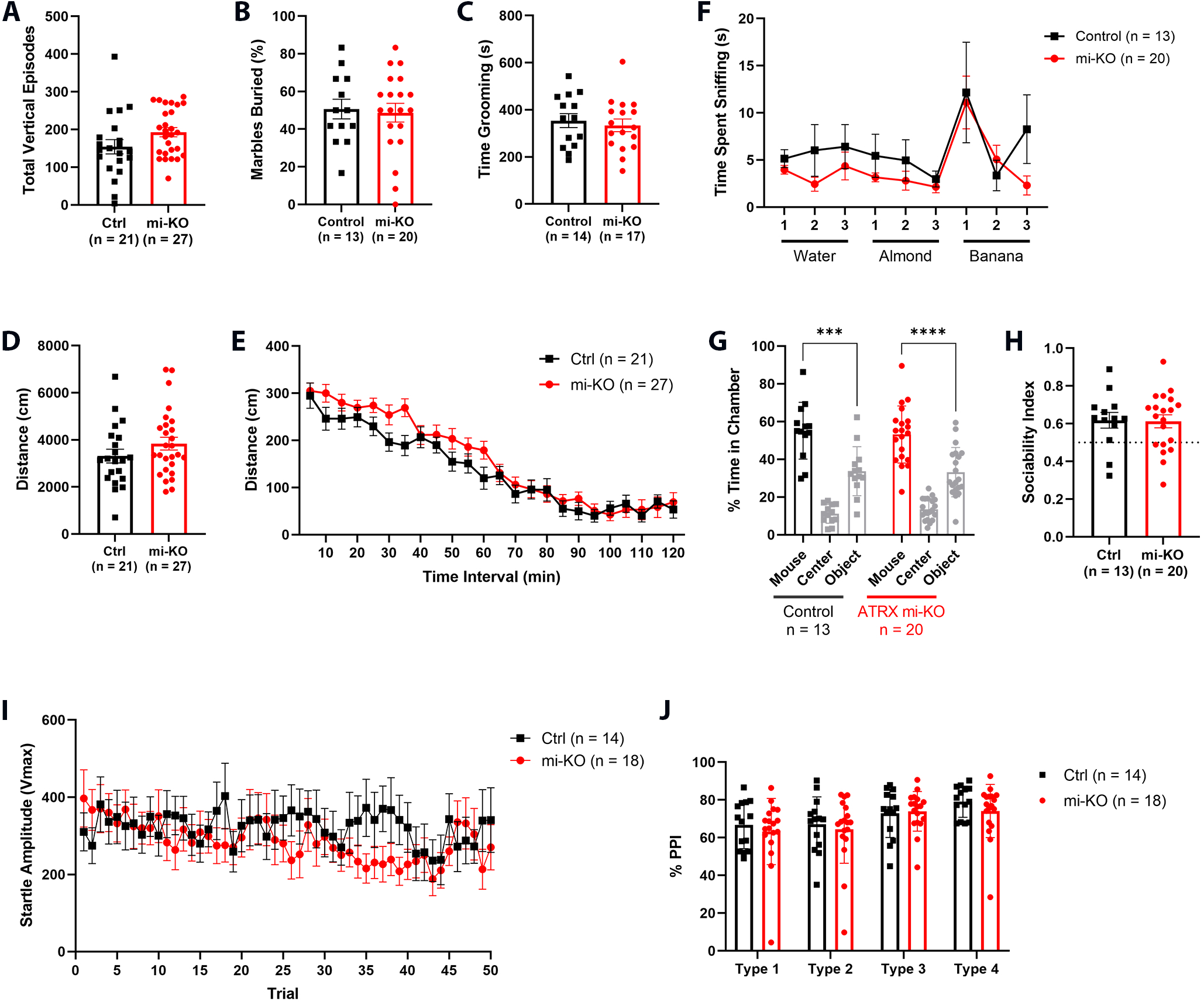
Normal activity levels and no repetitive behaviours, social deficits or impairments in sensory gating in mice upon perinatal ablation of ATRX in the microglia. **(a)** Percentage of marbles buried over 30 minutes during the marble burying test (t-test). **(b)** Time spent grooming over 30 minutes during the induced self-grooming test (t-test). **(c)** Number of vertical episodes over the 2-hour open field test (t-test). **(d)** Total distance travelled over 2 hours (t-test), and **(e)** distance travelled over 5-minute intervals in the open field test (two-way ANOVA). **(f)** Time spent sniffing a cotton swab saturated with either water, almond scent or banana scent over a 2-minute trial (two-way ANOVA). **(g)** Time spent in chambers containing a stranger mouse or a novel object, or in an empty chamber during the three-chamber sociability assay (two-way ANOVA). **(h)** Sociability index, calculated as the ratio of time spent in the chamber with the mouse to the total time spent in the chambers with the mouse and the object (t-test). **(i)** Amplitude of the startle response to 50 pulse-only trials (two-way ANOVA), and **(j)** pre-pulse inhibition for four different types of trials that differ in the time between pulses and the intensity of the pre-pulse stimulus, normalized to a baseline determined by the pulse-only trials (t-test). Type 1 = 30 ms, 75 dB; Type 2 = 30 ms, 80 dB; Type 3 = 100 ms, 75 dB, Type 4 = 100 ms, 80 dB. Ctrl: Cre+ control, mi-KO: ATRX microglia knockout. The number (n) of mice used for each genotype is indicated in parentheses. Data is presented as mean ± SEM. *** = P<0.001, **** = P<0.0001.

Given that olfactory deficits can confound social tasks, we first evaluated odour exploration [15], [16]. Over three 2-min trials with non-social and social odours, sniffing times did not differ between groups (F₁,₃₁ = 1.15, p = 0.29; **Fig. 6f**), confirming intact olfaction. In the three-chamber sociability assay both control and ATRX mi-KO mice spent significantly more time with a novel conspecific than with an empty cage or an inanimate object (mi-KO, F₂,₅₇ = 54.42, p < 0.0001; Ctrl, F₂,₃₆ = 44.96, p < 0.0001; **Fig. 6g**), and their sociability indices were virtually identical (p = 0.92; **Fig. 6h**). Finally, acoustic startle responses and prepulse inhibition (PPI) were examined to probe sensorimotor gating. The amplitude of the startle reflex across 50 consecutive 120-dB pulses was unaffected by genotype (F₁,₃₀ = 0.45, p = 0.51; **Fig. 6i**). Normalisation of startle magnitude to pulse-only trials revealed equivalent PPI at both prepulse intensities (75 dB, 80 dB) and intervals (100 ms, 110 ms) (all p > 0.25; **Fig. 6j**), indicating intact sensory gating.

Taken together, the absence of differences in stereotypy, locomotor activity, sociability, and PPI shows that perinatal ablation of ATRX in microglia does not produce autism-like behaviours in mice.

### The proportion of ATRX-null microglia decreases between 1 and 3 months of age

All behavioural assays described above were conducted when the mice were 3–6 months old, well after ATRX deletion had initially been confirmed at 1 month. To verify that the knockout persisted, we re-examined ATRX expression in adult brains. Coronal sections from three-month-old ATRX mi-KO mice and controls were co-stained for Iba1 and ATRX, and the percentage of ATRX-immunonegative microglia was quantified. In the cortex, the prevalence of ATRX-null microglia fell sharply: more than 90% of Iba1⁺ cells lacked ATRX at one month, whereas only ∼25% did so after three months (one-month: Ctrl n = 3, mi-KO n = 5; three-month: Ctrl n = 9, mi- KO n = 10; two-way ANOVA, genotype × age interaction F₁,₂₃ = 71.07, p = 0.0009; **Fig. 7a**). A smaller but significant reduction occurred in the hippocampus, where the fraction of ATRX-null microglia declined from ∼90% to ∼35% (F₁,₂₃ = 37.86, p = 0.0015; **Fig. 7b**) although there was some variability within the different brains. This marked drop suggests ongoing microglial turnover and replacement by newly generated, Cre-naïve cells. The decreased proportion of ATRX-null microglia in adulthood could potentially explain the absence of behaviour abnormalities in the 3–6-month-old cohort.

**Figure 7.**
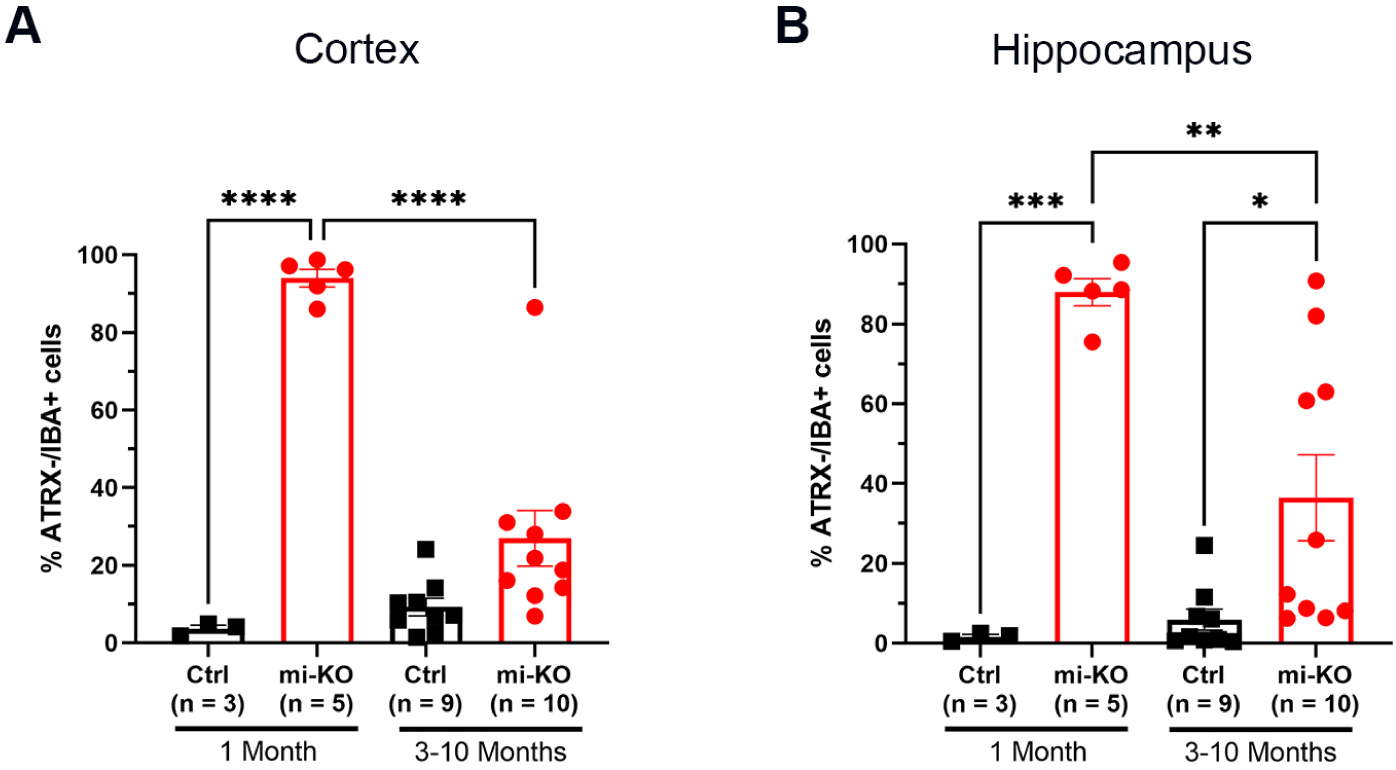
The number of ATRX KO microglia decreases over time in the cortex of the mi- KO mice. Quantification of brain cryosections from 1 month and 3-month-old mice immunostained with ATRX and the microglial marker IBA1, in **(a)** the cortex and **(b)** the hippocampus. Presented as the percentage of IBA1+ cells that are ATRX negative (two-way ANOVA). Ctrl: Cre+ control, mi-KO: ATRX microglia knockout. The number (n) of mice used for each genotype is indicated in parentheses. Data is presented as mean ± SEM. Multiple comparisons, * = P<0.05, ** = P<0.01, *** = P<0.001, ***** = P<0.0001.

## Discussion

This study set out to determine whether perinatal ablation of ATRX in microglia would cause more pronounced behavioral and cellular consequences than ATRX deletion in adult microglia, given the established role of microglia in early brain development and the severe cognitive deficits observed in ATR-X syndrome patients. While previous work has demonstrated that ATRX loss in adult microglia leads to cellular activation and memory impairment [13], our perinatal knockout model revealed a distinct outcome: ATRX-null microglia exhibited a robust reactive phenotype characterized by elevated CD68 expression and increased proliferation, but these changes did not translate into detectable deficits in anxiety, learning, memory, or autistic- like behaviors in adulthood.

It is important to highlight that, unlike a previous study reporting that Cre expression alone can induce a reactive phenotype in microglia [14], we observed no significant differences in the number of CD68-positive foci between Cre-positive and Cre-negative control mice. This suggests that the increased microglial reactivity observed in our model is not attributable to Cre expression itself. One likely explanation for this discrepancy lies in the method of tamoxifen administration: in our experiments, tamoxifen was delivered indirectly to pups through maternal milk, whereas the prior study administered tamoxifen directly to the pups via intragastric injection [14]. Delivery route might influence both the timing and magnitude of Cre activation, and a more gradual exposure through maternal milk may reduce the likelihood of acute microglial activation or off- target effects associated with direct dosing.

Despite increased proliferation, microglial density remained persistently reduced by approximately 20% at both one and three months following knockout. Although there was a trend toward elevated apoptosis, statistical significance was detected only in the CA2/3 region of the hippocampus, which alone does not fully account for the overall microglial loss. It is plausible that a slow, continuous rate of apoptosis, not readily detectable at isolated time points, cumulatively contributes to this reduction in cell density. Moreover, alternative cell death pathways may also play important roles in microglial attrition in this context. Pyroptosis, a pro-inflammatory form of programmed cell death characterized by plasma membrane rupture and the release of intracellular inflammatory mediators, could potentially contribute to microglial loss [25]. Similarly, cytorrhexis, which involves microglial dystrophy and fragmentation of the cytoplasm, may underlie microglial degeneration following prolonged activation [26].

Microglia participate in the modulation of myelination by engulfing excess OPCs, and abnormal myelination has been reported in ATR-X syndrome [17], [18], animal models of ASD as well as ASD patients [10], [12]. We detected no differences in OPC density or motor learning between ATRX mi-KO and control mice, in contrast to studies where reactive microglia induced OPC apoptosis *in vitro* [19]. This further supports the idea that compensatory mechanisms, possibly involving astrocytes or other glial cells, can maintain myelination and motor function even when microglia are dysfunctional.

The lack of behavioral changes in ATRX mi-KO mice differs from both the adult knockout model and from other studies that link reactive microglia to cognitive and social deficits [13], [20], [21], [22]. Additionally, post-mortem studies of patients with ASD have shown the presence of reactive microglia in brain tissue and cerebrospinal fluid [23], [24]. Several factors could contribute to the lack of behavioural deficits. First, the pattern of microglial activation we observed upon deletion of ATRX was transient. At one month of age, there was a significant increase in CD68-positive foci in microglia across all brain regions examined. By three months, this activation had partially resolved in the hippocampus but persisted in the cortex, even though the proportion of ATRX-null microglia in the cortex had decreased. The perinatal period is marked by high plasticity, which may allow for compensation that masks the effects of early microglial dysfunction. Studies have shown that even complete microglial ablation during development does not prevent normal synaptic maturation [25], implying that other glial populations, such as astrocytes, can take over essential roles in circuit refinement. Astrocytes have been shown to upregulate phagocytic machinery and compensate for microglial dysfunction or loss, supporting synaptic homeostasis when microglia are not fully functional [26]. A second potential reason for the absence of behaviour changes is that microglial ATRX ablation is temporally restricted to the perinatal period, after major developmental processes such as neurogenesis have largely concluded. Microglia emerge early in fetal brain development and play essential roles in regulating neural progenitor cell populations and synaptic pruning, making them particularly sensitive to immune challenges during this critical window. Finally, the timing of behavioral testing might also be a factor, as these assays were performed after the proportion of ATRX-null microglia had dropped, likely due to repopulation by wild-type cells. This timing may have reduced the ability to detect any behavioral effects stemming from early microglial perturbation.

In summary, perinatal ablation of ATRX in microglia causes persistent changes in microglial homeostasis and density but does not result in obvious behavioral deficits in adulthood, at least in the battery of tests included. These findings highlight the adaptability of the developing brain and the capacity for cellular compensation upon microglial dysfunction. While it refines our understanding of ATRX’s role in neurodevelopmental disorders, further mechanistic studies will be required to reveal the interplay between microglia, astrocytes, and other neural cell types in maintaining circuit homeostasis after early-life perturbations.

## Materials and Methods

### Animal Care and Husbandry

All animal procedures were conducted in accordance with the Animals for Research Act (Ontario, Canada) and approved by the Animal Care and Use Committee of the University of Western Ontario. Experimental design and reporting adhered to the ARRIVE guidelines. Mice were housed in a temperature- and humidity-controlled facility under a 12-hour light/12-hour dark cycle, with unrestricted access to standard chow and water. Female ATRX^loxp^ mice (C57BL/6 background) [27] were mated with male C57BL/6 mice expressing Cre^ER^ recombinase under the control of the *Cx3Cr1* promoter (B6.129P2(Cg)- Cx3cr1^tm2.1(cre/ERT2)Litt^/WganJ, MGI:5617710, RRID:ISMR JAX:021160). To enable visualization of Cre-mediated recombination, a subset of ATRX^loxp^ females were also bred with males carrying the tdTomato-Ai14 reporter allele (B6.Cg-Gt(ROSA)26Sor^tm14(CAG-tdTomato)Hze^/J, MGI:3809524, RRID:IMSR_JAX:007914) which drives robust tdTomato expression following Cre activity [28]. Conditional ablation of ATRX in microglia was achieved by administering tamoxifen to lactating dams, thereby delivering tamoxifen to pups via maternal milk. This approach induced ATRX deletion specifically in microglia of offspring harboring both ATRX^loxp^ and Cx3Cr1^CreER^ alleles (ATRX^loxp^;Cx3Cr1^CreER^). Littermates with either ATRX^loxp^ only (Cre-negative controls) or Cx3Cr1^CreER^ only (Cre-positive controls) served as expermental controls. Ear notch biopsies were collected from pups between postnatal days 7 and 12 (P7–P12) for PCR-based genotyping, as previously described [29]. Primer sequences used for genotyping are provided in Table 1.

**Table 1.**
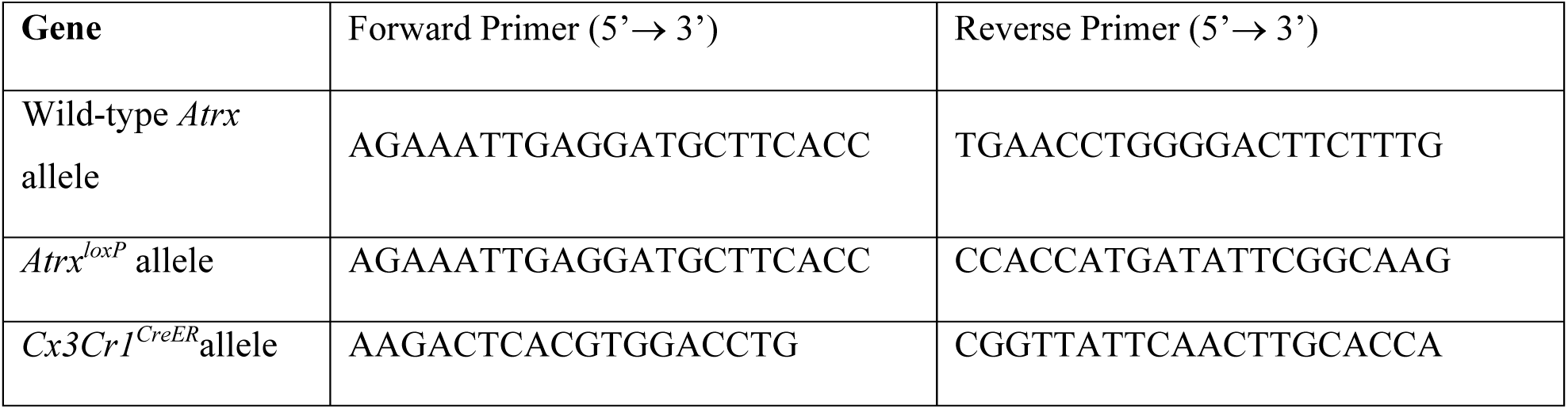
List of primers used for PCR genotyping.

### Tamoxifen administration

Tamoxifen (10 mg; Sigma Cat# T5648) was dissolved in 100µL 95% ethanol at 55°C for 10 minutes, followed by dilution with 900µL corn oil (Sigma Cat# C8267). Lactating dams were injected intraperitoneally with 1mg tamoxifen daily for 3 consecutive days, beginning when their pups were 2 days old (P2).

### Immunofluorescence

Mice were euthanized with CO_2_ at 1 month or 3 months of age and transcardially perfused with 25 mL 1x phosphate buffered saline (PBS) followed by 4% paraformaldehyde (PFA) in PBS. The brains were kept in 4% PFA overnight, and then washed 3 times with 1x PBS the following day before being placed in 30% sucrose/PBS solution. Brains were flash frozen in cryomatrix (Fisher Scientific Cat# 6769006), sectioned coronally to 10 µm in thickness (Leica CM 3050S) on Superfrost slides (Thermo Fisher Cat# 22-037-246) and stored at -80°C. For immunofluorescence staining, slides were removed from the freezer and post-fixed in 4% PFA for 15 minutes. Sections were rehydrated twice in 1x PBS for 5 minutes. For the IBA1, ATRX, Ki67, Casp3 and PDGFRα antibodies, antigen retrieval was performed by placing the slides in 10 mM sodium citrate at 95°C for 10 minutes. Slides were cooled, then washed 2 times for 5 minutes with 0.3% Triton X-100 in 1x PBS and incubated in blocking solution consisting of 5% donkey serum and 1% bovine serum albumin. Slides were incubated with primary antibody overnight at 4°C. The following day, slides were washed 3 times for 5 minutes with 0.3% Triton X-100 in 1x PBS and incubated with secondary antibody diluted in blocking solution for 1 hour at room temperature. Slides were wash twice with wash solution, counterstained with 1µg/mL DAPI (Millipore Sigma Cat# D9542) for 5 minutes and washed once more for 5 minutes before mounting with Permafluor (Thermo Fisher Cat# TA-006-FM). The following primary antibodies were used: anti-Iba1, rabbit polyclonal (1:800, FUJIFILM Wako Shibayagi Cat# 019-19741, RRID:AB_839504); anti-Iba1, goat polyclonal (1:300, Novus Cat# NB100-1028, RRID:AB_521594); anti-Iba1, chicken monoclonal (1:800, Synaptic Systems Cat #234 009, RRID:AB_2891282); anti-ATRX H-300, rabbit polyclonal (1:200, Santa Cruz Biotechnology Cat# sc-15408, RRID:AB_2061023); anti-CD68, rat monoclonal (1:200, Bio-Rad Cat# MCA1957, RRID:AB_322219); anti-Ki67, rabbit polyclonal (1:200, Abcam Cat# ab15580, RRID:AB_443209); anti-Cleaved Caspase-3, rabbit polyclonal (1:200, Cell Signaling Technology Cat# 9661, RRID:AB_2341188) anti-PDGFRα, rabbit monoclonal (1:200, Abcam Cat# ab203491, RRID:AB_2892065). The secondary antibodies used were goat anti-rabbit-Alexa Fluor 594 (1:800, Thermo Fisher Scientific Cat# A-11012, RRID:AB_2534079), donkey anti-rabbit-Alexa Fluor 488 (1:800, Thermo Fisher Scientific Cat# A-21206, RRID:AB_2535792), goat anti-chicken-Alexa Fluor 488 (1:800, Thermo Fisher Scientific Cat# A-11039, RRID:AB_2534096), donkey anti-goat-Alexa Fluor 594 (1:800, Thermo Fisher Scientific Cat# A-11058, RRID:AB_2534105), goat anti-rat-Alexa Fluor 488 (1:800, Thermo Fisher Scientific Cat# A- 11006, RRID:AB_2534074), and goat anti-rabbit-Alexa Fluor 647 (1:800, Thermo Fisher Scientific Cat# A-21244, RRID:AB_2535812).

### Microscopy and Image Analysis

Immunofluorescence images were acquired using a Leica DM6000 B inverted fluorescence microscope equipped with a Hamamatsu ORCA-ER-1394 digital camera. Image acquisition was performed with Volocity Acquisition software (RRID:SCR_002668), and subsequent image processing and adjustments were carried out using Volocity (Demo Version 6.0.1) and Adobe Photoshop (Version 20.0.4, RRID:SCR_014199). All adjustments were applied equally across experimental groups and were limited to linear changes in brightness and contrast. Regions of interest included the CA1, CA2, CA3, and dentate gyrus subfields of the hippocampus, as well as the cortical area immediately above the CA2 region in both hemispheres. For each experimental condition, a minimum of three biological replicates (n ≥ 3 mice per group) were analyzed. Quantification was performed on 4–5 anatomically matched coronal brain sections per animal. Cell counts and fluorescence intensity measurements were conducted using ImageJ software (Version 1.53a, RRID:SCR_003070). All quantification was performed by experimenters blinded to genotype and experimental condition. Data were averaged across sections for each animal prior to statistical analysis.

### Behavioral Testing

Behavioral assessments were conducted between 9:00 AM and 4:00 PM on a minimum of 11 animals per genotype. Mice were randomly assigned to experimental groups, and all behavioral data were collected and analyzed by experimenters blinded to genotype. Mouse performance in behavioral tasks was quantified using automated, software-based scoring systems.

### Open Field Test

The open field test was performed as previously described [30], [31]. Mice were acclimated to the testing room for 30 minutes prior to the start of the experiment to minimize handling and transport stress. Each mouse was individually placed in the center of a square open field arena (20 cm × 20 cm × 30 cm; width × length × height) and allowed to explore freely for a total of 2 hours. Locomotor activity was automatically recorded in 5-minute intervals using Fusion software (AccuScan Instruments). The arena was thoroughly cleaned with 70% ethanol between trials to eliminate olfactory cues.

### Light Dark Box

The light dark box test was performed based on the original protocol by Crawley et al. [32], with the following modifications. Mice were habituated to the testing room for 30 minutes prior to testing. The apparatus consisted of a square chamber divided into two equal-sized compartments: one illuminated (light) and one covered (dark). At the start of each trial, mice were placed in the center of the light compartment and allowed to explore both compartments freely for 15 minutes. Locomotor activity and time spent in each compartment were recorded in 5-minute intervals using Fusion software (AccuScan Instruments). The apparatus was cleaned with 70% ethanol between trials to prevent odor-based confounds.

### Elevated Plus Maze

The elevated plus maze test was conducted as previously described [30]., with minor modifications. Mice were habituated to the testing room for 30 minutes prior to the experiment to minimize stress. The apparatus consisted of two opposing open arms and two opposing closed arms (each measuring 38 cm from the center by 7 cm). Mice were placed at the center of the maze facing an open arm and allowed to explore freely for 5 minutes. Video tracking and analysis were performed using ANY-maze software (Stoelting Co.), which automatically recorded the time spent in open and closed arms, as well as the number of entries into each arm. The animal’s center point was used to determine zone entries.

### Marble Burying Assay

The marble burying assay was performed as previously described [33], [34]. Mice were acclimated to the testing room for 30 minutes prior to testing. Each mouse was placed in a standard cage filled with 5 cm of bedding, with 12 marbles spaced out evenly on top of the bedding. Mice were allowed to explore and interact with the marbles for 30 minutes. At the end of the trial, mice were gently removed from the cage without disturbing the bedding. The number of marbles buried at least two-thirds (2/3) beneath the bedding was counted by an experimenter blinded to genotype. The cage and marbles were cleaned with 70% ethanol between trials to prevent odor-based confounds.

### Induced Grooming Test

The induced self-grooming test was performed as previously described [33], [35] with modifications [33]. Mice were brought into the room 30 minutes prior to the start of the test for habituation. Mice were misted with water on their back using a spray bottle held approximately 10 cm away from the mouse and then placed in an empty cage for 30 minutes. Videos were captured using the ANY-maze video tracking software, and the time spent grooming by each mouse over 30 minutes was manually recorded.

### Olfaction

Testing for odor discrimination was performed as previously described to assess olfaction in the mice [33], [36]. Mice were habituated in an empty cage with a wire lid for 30 minutes prior to the start of the test in a separate room. A cotton swab saturated with either water, almond or banana odour was positioned through the water bottle opening of the cage wire. Each mouse was exposed to the scent for three consecutive trials for each odour, each lasting 2 minutes. The amount of time spent sniffing the cotton swab was recorded manually.

### Three-Chamber Sociability

The social preference test was performed as previously described [33], [37], [38]. Mice were acclimated to the testing room for 30 minutes prior to the start of the experiment to reduce stress and novelty effects. The apparatus consisted of a rectangular chamber divided into three equal compartments, each connected by small openings that allowed free movement between chambers. During the initial habituation phase, the test mouse was placed in the center chamber and allowed to freely explore all three chambers for 10 minutes. Following habituation, the mouse was temporarily removed from the apparatus. A novel object was placed in one of the side chambers, while an unfamiliar, age- and sex- matched stranger mouse was confined within an identical wire cage in the opposite side chamber. The location of the novel object and stranger mouse was counterbalanced across subjects to avoid side bias. The test mouse was then reintroduced into the center chamber and allowed to explore the entire apparatus freely for an additional 10 minutes. Behavior was recorded using ANY-maze video tracking software which automatically measured the time spent in each chamber. The sociability index was calculated as the ratio of time spent in the chamber containing the stranger mouse to the total time spent in the chambers containing either the stranger mouse or the novel object.

### Prepulse Inhibition of the Startle Response

The pre-pulse inhibition and startle response tests were performed as previously described with slight modifications [33], [39]. All testing was conducted using the SR-LAB Startle Response System (San Diego Instruments). Mice were habituated to the testing apparatus for two consecutive days prior to testing. During each habituation session, animals were placed individually in the startle chamber and exposed to a continuous background noise of 68 dB for 5 minutes. On the test day, mice were placed in the chamber and acclimated for 10 minutes with 68 dB background noise. The test session consisted of two phases. For the acoustic startle response (ASR) phase, mice received 50 startle pulses (115 dB, 20 ms duration) presented at 20-second intervals. The magnitude of the startle response, measured as the peak movement of the animal within the chamber, was automatically recorded by the SR-LAB software. Following the ASR phase, mice were exposed to 10 sets of 5 trials consisting of a prepulse varying in intensity (75 dB or 80 dB) and latency to the acoustic startle (30 ms or 100 ms), and one startle only trial. The startle response was recorded by the software and normalized to the startle-only trial. The average response for each of the different trial types was calculated and normalized to the startle trial alone.

### Y-Maze

The Y-maze was performed as previously described with slight modifications [30], [40]. Mice were acclimated to the testing room for 30 minutes prior to the start of the experiment to minimize stress and novelty effects. Each mouse was then placed at the end of one arm of a symmetrical Y-shaped maze (each arm measuring 38 cm from the center by 6 cm) and allowed to explore freely for a 5-minute trial. Mouse behavior was recorded using ANY-maze video tracking software, which automatically tracked arm entries and movement patterns. Spontaneous alternation was defined as consecutive entries into all three arms without repeating an arm.

### Morris Water Maze

The Morris water maze was performed as previously described with slight modifications [30], [41]. Mice were placed in individual cages to habituate for 10 minutes prior to the start of the test. The maze consisted of a circular pool (diameter: 1.5 m) filled with water maintained at 25–27°C. Extra-maze visual cues were positioned on the walls surrounding each quadrant to facilitate spatial navigation. An escape platform (diameter: 10 cm) was submerged 1.5 cm below the water surface in a fixed location within one quadrant. From days 1 to 4, mice underwent spatial learning trials. Each mouse received four training trials per day, with a minimum inter-trial interval of 15 minutes. For each trial, mice were released from one of four randomized start positions and allowed up to 60 seconds to locate the hidden platform. If a mouse failed to find the platform within 60 seconds, it was gently guided to the platform and allowed to remain there for 10 seconds before being returned to its cage. On days 5 and 12, probe trials were conducted to assess spatial memory retention. The platform was removed, and each mouse was placed in the pool for a single 60-second trial. The time spent in each quadrant, and the swimming distance and swimming speed were recorded by the ANY-maze video tracking system.

### Contextual Fear Conditioning

The contextual fear paradigm was performed as previously described, with some modifications [30]. Mice were acclimated to the testing room for 30 minutes prior to the start of the experiment to minimize stress. The conditioning chamber consisted of a metal grid floor (20 cm × 10 cm) enclosed by walls featuring distinct visual cues on opposing sides to facilitate context discrimination. The chamber was thoroughly cleaned with 70% ethanol between trials to eliminate olfactory cues. On the training day, each mouse was placed in the chamber and allowed to freely explore for 180 seconds. After 150 seconds, a single foot shock (2 mA, 180 V, 2 seconds) was delivered through the metal grid floor. Mouse behavior was recorded using ANY-maze video tracking software, which continuously monitored locomotion and activity. Twenty-four hours after training, mice were returned to the same chamber for a 600-second (10- minute) test session. No shocks were administered during this session. Freezing behavior, defined as the absence of all movement for at least 850 ms, was automatically detected and quantified by ANY-maze software. The total duration of freezing was used as an index of contextual fear memory.

### Rotarod

The rotarod test was performed as previously described with modifications [42]. Mice were brought into the room 30 minutes prior to the start of the test for habituation. Mice were placed on the Rotarod apparatus. The rotating speed was gradually increased from 5 rpm to 35 rpm. The latency to fall off the rotarod, up to a maximum of 6 minutes, was recorded automatically by sensors on the apparatus. On the first day, 10 trials were performed, with at least 10 minutes rest in between trials. 24 hours later, another 4 trials were completed.

### Statistical Analysis

All data was analyzed using GraphPad Prism software. Student’s *t-test* (unpaired, two-tailed) was performed to compare two data sets, while multiple data sets were compared using one- or two-way ANOVA with post hoc Tukey test for multiple comparisons. Data is presented as the mean ± SEM and p values less than 0.05 were considered significant.

## Data Availability

All data generated or analyzed in this study are included in this published article.

## Acknowledgements

We are grateful to Drs. Vania Prado and Marco Prado for the Cx3Cr1^ER^ mice, and for access to the Robarts Research Institute neurobehavioral core facility along with the guidance of the facility manager Matthew Cowan.

## Funding Declaration

This work was supported by the Canadian Institutes for Health Research operating grant to N.G.B. (FRN#183661) and by BrainsCAN through the Canada First Research Excellence Fund. K.M. received a Summer Studentship and a Graduate Studentship from the Department of Paediatrics at Western University, as well as an Ontario Graduate Scholarship and the C. Allison Kingsley Research Grant from the Department of Psychiatry at Western University. S.S. received a Children’s Health Research Institute Trainee award funded by the Children’s Health Foundation.

## Author Contribution

K.Y.M. contributed to conceptualization, data curation, formal analysis, and writing - original draft and review & editing; S.S contributed to conceptualization and data curation. N.G.B contributed to conceptualization, funding acquisition, supervision, writing original draft and review & editing.

## Additional Information

The authors declare no competing interests.

## Notes

### Competing Interest Statement

The authors have declared no competing interest.

## References

[1] D. J. Picketts, D. R. Higgs, S. Bachoo, D. J. Blake, O. W. Quarrell, and R. J. Gibbons, ‘ATRX encodes a novel member of the SNF2 family of proteins: mutations point to a common mechanism underlying the ATR-X syndrome.’, Hum Mol Genet, vol. 5, no. 12, pp. 1899–1907, Dec. 1996, doi: 10.1093/hmg/5.12.1899.

[2] X. Gong et al., ‘Analysis of X chromosome inactivation in autism spectrum disorders’, Am J Med Genet B Neuropsychiatr Genet, vol. 147B, no. 6, pp. 830–835, Sep. 2008, doi: 10.1002/ajmg.b.30688.

[3] R. J. Gibbons, D. J. Picketts, L. Villard, and D. R. Higgs, ‘Mutations in a putative global transcriptional regulator cause X-linked mental retardation with alpha-thalassemia (ATR- X syndrome).’, Cell, vol. 80, no. 6, pp. 837–845, Mar. 1995, doi: 10.1016/0092-8674(95)90287-2.

[4] C. Badens et al., ‘ATRX syndrome in a girl with a heterozygous mutation in the ATRX Zn finger domain and a totally skewed X-inactivation pattern’, Am J Med Genet A, vol. 140A, no. 20, pp. 2212–2215, Oct. 2006, doi: 10.1002/ajmg.a.31400.

[5] U.-K. Hanisch and H. Kettenmann, ‘Microglia: active sensor and versatile effector cells in the normal and pathologic brain’, Nat Neurosci, vol. 10, no. 11, pp. 1387–1394, 2007, doi: 10.1038/nn1997.

[6] R. C. Paolicelli et al., ‘Microglia states and nomenclature: A field at its crossroads’, Neuron, vol. 110, no. 21, pp. 3458–3483, Nov. 2022, doi: 10.1016/j.neuron.2022.10.020.

[7] H. Kettenmann, U.-K. Hanisch, M. Noda, and A. Verkhratsky, ‘Physiology of Microglia’, Physiol Rev, vol. 91, no. 2, pp. 461–553, Apr. 2011, doi: 10.1152/physrev.00011.2010.

[8] Y. Fujita and T. Yamashita, ‘Neuroprotective function of microglia in the developing brain’, Neuronal Signal, vol. 5, no. 1, p. NS20200024, Jan. 2021, doi: 10.1042/NS20200024.

[9] J. L. Frost and D. P. Schafer, ‘Microglia: Architects of the Developing Nervous System’, Trends Cell Biol, vol. 26, no. 8, pp. 587–597, Aug. 2016, doi: 10.1016/j.tcb.2016.02.006.

[10] A. D. Nemes-Baran, D. R. White, and T. M. DeSilva, ‘Fractalkine-Dependent Microglial Pruning of Viable Oligodendrocyte Progenitor Cells Regulates Myelination’, Cell Rep, vol. 32, no. 7, p. 108047, Aug. 2020, doi: 10.1016/j.celrep.2020.108047.

[11] R. C. Paolicelli et al., ‘Synaptic pruning by microglia is necessary for normal brain development.’, Science, vol. 333, no. 6048, pp. 1456–1458, Sep. 2011, doi: 10.1126/science.1202529.

[12] M. Graciarena, A. Seiffe, B. Nait-Oumesmar, and A. M. Depino, ‘Hypomyelination and Oligodendroglial Alterations in a Mouse Model of Autism Spectrum Disorder.’, Front Cell Neurosci, vol. 12, p. 517, 2018, doi: 10.3389/fncel.2018.00517.

[13] S. Shafiq, et al., ‘Viral mimicry and memory deficits upon microglial deletion of ATRX’, *bioRxiv*, p. 2024.05.07.592875, Jan. 2024, doi: 10.1101/2024.05.07.592875.

[14] V. Sahasrabuddhe and H. S. Ghosh, ‘Cx3Cr1-Cre induction leads to microglial activation and IFN-1 signaling caused by DNA damage in early postnatal brain’, Cell Rep, vol. 38, no. 3, Jan. 2022, doi: 10.1016/j.celrep.2021.110252.

[15] B. C. Ryan, N. B. Young, S. S. Moy, and J. N. Crawley, ‘Olfactory cues are sufficient to elicit social approach behaviors but not social transmission of food preference in C57BL/6J mice’, Behavioural Brain Research, vol. 193, no. 2, pp. 235–242, Nov. 2008, doi: 10.1016/J.BBR.2008.06.002.

[16] S. H. de la Zerda et al., ‘Social recognition in laboratory mice requires integration of behaviorally-induced somatosensory, auditory and olfactory cues’, Psychoneuroendocrinology, vol. 143, p. 105859, Sep. 2022, doi: 10.1016/J.PSYNEUEN.2022.105859.

[17] J. S. Lee et al., ‘Alpha-thalassemia X-linked intellectual disability syndrome identified by whole exome sequencing in two boys with white matter changes and developmental retardation’, Gene, vol. 569, no. 2, pp. 318–322, 2015, doi: 10.1016/j.gene.2015.04.075.

[18] T. Wada et al., ‘Neuroradiologic Features in X-linked α-Thalassemia/Mental Retardation Syndrome’, American Journal of Neuroradiology, vol. 34, no. 10, p. 2034, Oct. 2013, doi: 10.3174/ajnr.A3560.

[19] Y. Li et al., ‘Microglia activation triggers oligodendrocyte precursor cells apoptosis via HSP60’, Mol Med Rep, vol. 16, no. 1, pp. 603–608, 2017.

[20] H. Jung et al., ‘LPS induces microglial activation and GABAergic synaptic deficits in the hippocampus accompanied by prolonged cognitive impairment’, Sci Rep, vol. 13, no. 1, p. 6547, 2023, doi: 10.1038/s41598-023-32798-9.

[21] R. Chesworth et al., ‘Spatial Memory and Microglia Activation in a Mouse Model of Chronic Neuroinflammation and the Anti-inflammatory Effects of Apigenin’, Front Neurosci, vol. 15, 2021, [Online]. Available: https://www.frontiersin.org/articles/10.3389/fnins.2021.699329

[22] M. Wadhwa et al., ‘Inhibiting the microglia activation improves the spatial memory and adult neurogenesis in rat hippocampus during 48 h of sleep deprivation’, J Neuroinflammation, vol. 14, no. 1, p. 222, 2017, doi: 10.1186/s12974-017-0998-z.

[23] J. T. Morgan et al., ‘Microglial activation and increased microglial density observed in the dorsolateral prefrontal cortex in autism’, Biol Psychiatry, vol. 68, no. 4, pp. 368–376, 2010.

[24] D. L. Vargas, C. Nascimbene, C. Krishnan, A. W. Zimmerman, and C. A. Pardo, ‘Neuroglial activation and neuroinflammation in the brain of patients with autism’, Ann Neurol, vol. 57, no. 1, pp. 67–81, Jan. 2005, doi: 10.1002/ana.20315.

[25] M. O’Keeffe et al., ‘Typical development of synaptic and neuronal properties can proceed without microglia in the cortex and thalamus’, Nat Neurosci, vol. 28, no. 2, pp. 268–279, 2025, doi: 10.1038/s41593-024-01833-x.

[26] H. Konishi et al., ‘Astrocytic phagocytosis is a compensatory mechanism for microglial dysfunction’, EMBO J, vol. 39, no. 22, p. e104464, Nov. 2020, doi: 10.15252/embj.2020104464.

[27] N. G. Bérubé et al., ‘The chromatin-remodeling protein ATRX is critical for neuronal survival during corticogenesis’, J Clin Invest, vol. 115, no. 2, pp. 258–267, Feb. 2005, doi: 10.1172/JCI22329.

[28] L. Madisen et al., ‘A robust and high-throughput Cre reporting and characterization system for the whole mouse brain’, Nat Neurosci, vol. 13, no. 1, pp. 133–140, 2010, doi: 10.1038/nn.2467.

[29] M. A. Pena-Ortiz, S. Shafiq, M. E. Rowland, and N. G. Bérubé, ‘Selective isolation of mouse glial nuclei optimized for reliable downstream omics analyses’, J Neurosci Methods, vol. 369, p. 109480, 2022, doi: 10.1016/j.jneumeth.2022.109480.

[30] R. J. Tamming, J. R. Siu, Y. Jiang, M. A. M. Prado, F. Beier, and N. G. Bérubé, ‘Mosaic expression of Atrx in the mouse central nervous system causes memory deficits’, Dis Model Mech, vol. 10, no. 2, pp. 119–126, Feb. 2017, doi: 10.1242/dmm.027482.

[31] A. C. Martyn et al., ‘Elimination of the vesicular acetylcholine transporter in the forebrain causes hyperactivity and deficits in spatial memory and long-term potentiation’, Proceedings of the National Academy of Sciences, vol. 109, no. 43, pp. 17651–17656, Oct. 2012, doi: 10.1073/pnas.1215381109.

[32] J. Crawley and F. K. Goodwin, ‘Preliminary report of a simple animal behavior model for the anxiolytic effects of benzodiazepines.’, Pharmacol Biochem Behav, vol. 13, no. 2, pp. 167–70, Aug. 1980, doi: 10.1016/0091-3057(80)90067-2.

[33] N. Martin-Kenny and N. G. Bérubé, ‘Effects of a postnatal Atrx conditional knockout in neurons on autism-like behaviours in male and female mice.’, J Neurodev Disord, vol. 12, no. 1, p. 17, Jun. 2020, doi: 10.1186/s11689-020-09319-0.

[34] R. M. J. Deacon, ‘Digging and marble burying in mice: simple methods for in vivo identification of biological impacts’, Nat Protoc, vol. 1, no. 1, pp. 122–124, 2006, doi: 10.1038/nprot.2006.20.

[35] A. V Kalueff, J. W. Aldridge, J. L. LaPorte, D. L. Murphy, and P. Tuohimaa, ‘Analyzing grooming microstructure in neurobehavioral experiments.’, Nat Protoc, vol. 2, no. 10, pp. 2538–44, 2007, doi: 10.1038/nprot.2007.367.

[36] E. P. Arbuckle, G. D. Smith, M. C. Gomez, and J. N. Lugo, ‘Testing for odor discrimination and habituation in mice.’, J Vis Exp, no. 99, p. e52615, May 2015, doi: 10.3791/52615.

[37] S. Jamain et al., ‘Reduced social interaction and ultrasonic communication in a mouse model of monogenic heritable autism’, Proc Natl Acad Sci U S A, vol. 105, no. 5, pp. 1710–1715, Feb. 2008, doi: 10.1073/pnas.0711555105.

[38] A. El-Kordi et al., ‘Development of an autism severity score for mice using Nlgn4 null mutants as a construct-valid model of heritable monogenic autism’, Behavioural Brain Research, vol. 251, pp. 41–49, 2013, doi: 10.1016/j.bbr.2012.11.016.

[39] B. Valsamis and S. Schmid, ‘Habituation and prepulse inhibition of acoustic startle in rodents.’, J Vis Exp, no. 55, p. e3446, Sep. 2011, doi: 10.3791/3446.

[40] B. M. De Castro et al., ‘Reduced expression of the vesicular acetylcholine transporter causes learning deficits in mice’, Genes Brain Behav, vol. 8, no. 1, pp. 23–35, Feb. 2009, doi: 10.1111/j.1601-183X.2008.00439.x.

[41] C. V Vorhees and M. T. Williams, ‘Morris water maze: procedures for assessing spatial and related forms of learning and memory.’, Nat Protoc, vol. 1, no. 2, pp. 848–58, 2006, doi: 10.1038/nprot.2006.116.

[42] M. P. McFadyen, G. Kusek, V. J. Bolivar, and L. Flaherty, ‘Differences among eight inbred strains of mice in motor ability and motor learning on a rotorod’, Genes Brain Behav, vol. 2, no. 4, pp. 214–219, Aug. 2003, doi: 10.1034/j.1601-183X.2003.00028.x.

